# Human iPSC-CMs and in-silico technologies define mechanisms and accelerate targeted pharmacogenetics in hypertrophic cardiomyopathy

**DOI:** 10.1101/2022.06.08.495324

**Authors:** Francesca Margara, Yiangos Psaras, Zhinuo Jenny Wang, Manuel Schmid, Ruben Doste, Amanda Garfinkel, Giuliana G. Repetti, Jonathan Seidman, Christine Seidman, Blanca Rodriguez, Christopher N. Toepfer, Alfonso Bueno-Orovio

## Abstract

Cardiomyopathies have unresolved genotype-phenotype relationships and lack disease-specific treatments. Here we identify genotype-specific pathomechanisms and therapeutic targets combining experimental hiPSC-CM modelling and human-based cardiac electromechanical in-silico modelling and simulation bridging from specific mutations to clinical biomarkers. We select hypertrophic cardiomyopathy as a challenge for this approach and study genetic variations that mutate proteins of the thick (*MYH7*^R403Q/+^) and thin filaments (*TNNT2*^R92Q/+^, *TNNI3*^R21C/+^) of the cardiac sarcomere. We show that destabilisation of myosin super relaxation drives disease in *MYH7*^R403Q/+^ with secondary effects on thin filament activation, which are corrected by Mavacamten. Thin filament variants *TNNT2*^R92Q/+^ and *TNNI3*^R21C/+^ share calcium regulation-related pathomechanisms, for which Mavacamten provides incomplete salvage. We define the ideal characteristics of a novel thin filament-targeting compound and show its efficacy in-silico. We demonstrate that hybrid human-based hiPSC-CM and in-silico studies accelerate pathomechanism discovery and classification testing, improving clinical interpretation of genetic variants, and directing rational therapeutic targeting and design.

## Introduction

Genetic diseases lack targeted and disease-specific treatment options that therapeutically address causal disease mechanisms^1^. This is complicated by unresolved disease pathophysiology and population heterogeneity. Human-based modelling and simulation studies address these challenges by uncovering mechanisms bridging from mutation to clinical biomarkers, in combination with experimental and clinical data^2^. Thus, they can unravel insights into key features of disease pathomechanisms, improving therapeutic development and supporting precision medicine in the clinic.

Among genetic diseases, hypertrophic cardiomyopathy (HCM) is a common condition. Affecting approximately 1 in 500 people, HCM can cause arrhythmias, heart failure, and sudden cardiac death in affected individuals^3^. HCM is caused predominantly by mutations in sarcomere genes^4^, most abundantly in *MYH7, MYBP3, TNNT2*, and *TNNI3*. Within each of these genes there are many possible causative variants of HCM, with diverse pathomechanisms. Establishing genotype-phenotype relationships and pathomechanisms in HCM is the main bottleneck to clinical translation and precision medicine. This is even with the implementation of novel technologies such as CRISPR/Cas-9, human induced pluripotent stem cell-derived cardiomyocytes (hiPSC-CMs), and cutting-edge protein expression systems, which have allowed more rapid phenotyping.

Most HCM variants fall within genes that encode proteins of the cardiac sarcomere, which can be robustly modelled in-silico^5^. Human in-silico models enable simulation of force generation by calcium regulation of the troponin complex and the cyclical interactions between myosin and actin^6^. This is important as altered force at the myofilament can drive pathogenesis of many forms of cardiomyopathy, and novel therapeutics aim to modulate cardiac contractility for treating these therapeutically orphan heart diseases^7^. Human-based in-silico frameworks twinned with hiPSC-CM data can be leveraged to simulate single cell function and scale to entire chambers of the heart^8^ to accelerate phenotyping and drug discovery.

We therefore use a human in-silico modelling and simulation framework integrated with CRISPR/Cas-9 and hiPSC-CM modelling of disease, and clinical data, to accelerate biological and clinical investigation (Fig. 1). This hybrid human-based framework defines novel disease pathomechanisms across diverse HCM-causative genetic variants. We use this information to predict and explain divergent efficacy of Mavacamten9^,10^ (now commercially marketed as Camzyos), a first-in-class HCM-targeted small molecule, across HCM genotypes. We provide digital evidence of mechanistic pharmacological targets that can be used to tailor genotype-specific phenotype correction, as validated with hiPSC-CMs.

**Fig. 1.**
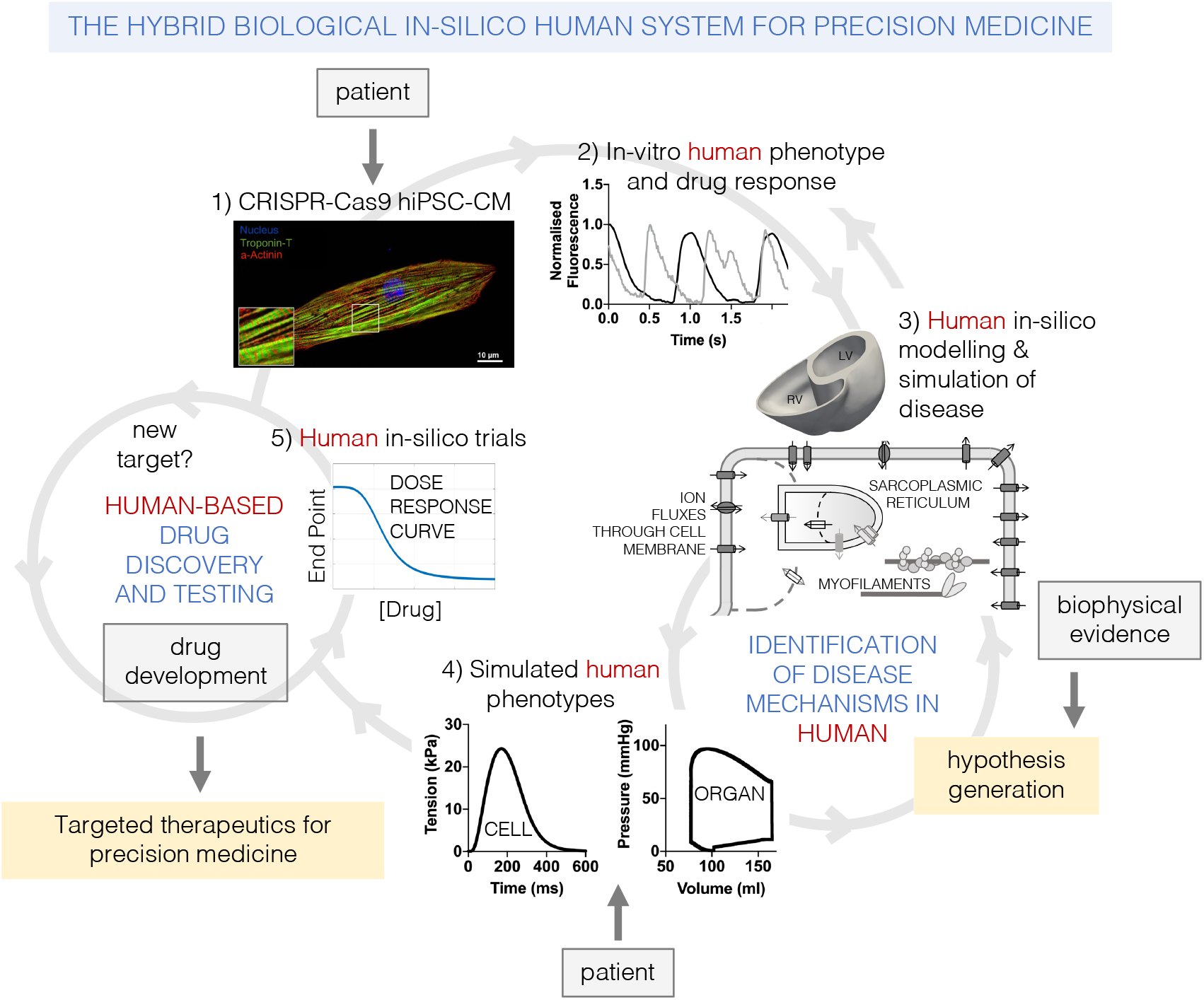
The hybrid biological and in-silico human system defined in this work to synergistically accelerate hypothesis generation and testing in variant classification, pathomechanism discovery, and therapeutic targeting. From patient genotype, hiPSC-CMs can be generated via CRISPR-Cas9 to express patient’s genetic variant (1). Cellular phenotypes and responses to pharmaceutical interventions can be studied in-vitro (2). Based on this and available biophysical evidence on mutational protein function, hypotheses on variant-specific disease mechanisms can be formulated. These are tested in in-silico human models of cardiac electromechanical function, from cell to organ (3). In-silico human disease models, informed by clinical data, enable simulation of the multiscale features of disease phenotypes (4). Simulated phenotypes are compared to in-vitro and clinical data to test whether hypothesised disease mechanisms explain observed phenotypes. Once a phenotype is correctly accounted for, in-silico drug trials can be conducted to test the safety and efficacy of pharmacological therapies (5). Based on in-silico evaluations, drugs can then proceed into further drug testing stages. In-silico trials can also identify new therapeutic targets and provide evidence to inform the best targeted therapeutic approach for the individual patient.

We propose this hybrid human-based framework can be extended widely within cardiovascular medicine to accelerate precision medicine.

## Results

### Simulating human HCM defines key mechanistic differences between *MYH7, TNNT2*, and *TNNI3* variants

Similar contractile profiles are observed in hiPSC-CMs expressing different HCM genetic variants across the *MYH7* (β-myosin heavy chain), *TNNT2* (troponin T), and *TNNI3* (troponin I) genes. A phenotype of hypercontractility and slowed relaxation is reported for the *MYH7*^R403Q/+^, *TNNT2*^R92Q/+^, and *TNNI3*^R21C/+^ variants^11-14^, which occurs alongside prolonged calcium decay times, and a faster time to transient peak observed in *TNNI3*^R21C/+ 11,12,14^.

We conducted human simulation studies to determine the mechanistic pathways that explain these similar cellular phenotypes, considering their diverse clinical manifestations. We generated populations of in-silico virtual cardiomyocytes using our extended cellular electromechanical model of human cardiomyocyte electromechanical function^15^ (Supplementary Figs. 1 and 2). Within this model we integrated variant-specific biophysical evidence of pathogenesis.

### MYH7^R403Q/+^ decreases myosin SRX increasing thin filament activation causing cellular HCM

We generated in-silico models of *MYH7*^R403Q/+^ cardiomyocytes with experimentally-informed^13^ reduced myosin super relaxation (SRX) (Fig. 2a,b). The myosin SRX deficit results in higher tension amplitude than control cells in-silico (Fig. 2c,d), corroborated by increased sarcomere shortening in *MYH7*^R403Q/+^ hiPSC-CMs^13^ (Fig. 2e). Reduced myosin SRX increases the disordered relaxed (DRX) myosin conformation, which becomes available to form crossbridges and drives hypercontractility^13,16^.

**Fig. 2.**
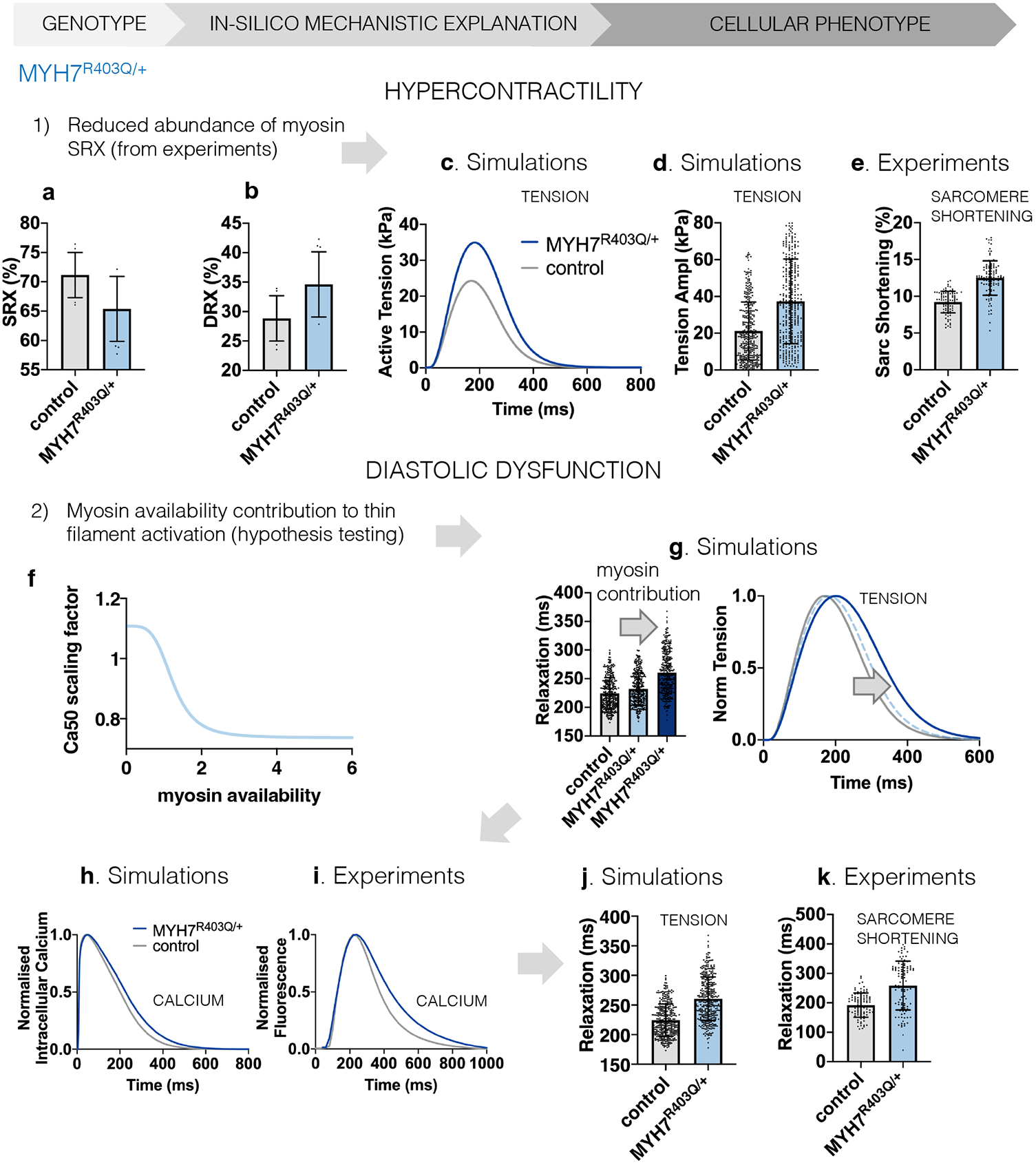
Mechanistic explanation of the MYH7^R403Q/+^ cellular phenotype of hypercontractility and diastolic dysfunction through human-based modelling and simulation informed by biophysical evidence. **a,b**, Experimental evidence on the mutation-induced change in myosin conformations leading to larger myosin availability under MYH7^R403Q/+^ used to inform modelling and simulation. **c**, Comparison of simulated active tension waveforms in control and under MYH7^R403Q/+^. **d,e**, Simulated MYH7^R403Q/+^ cardiomyocytes with larger myosin availability develop higher tension amplitude compared to control (**d**) in line with experimental hiPSC-CM data (**e**). **f**, Hypothesis tested in simulations to explain the pathway behind impaired relaxation in MYH7^R403Q/+^, i.e. the myosin-based contribution to thin filament activation. The scaling factor of Ca50 is reported. Ca50 represents the calcium concentration at half maximal thin filament activation and has μM unit. Therefore, a larger Ca50 value means lower calcium sensitivity and vice versa. **g**, Effect of myosin contribution to thin filament activation on simulated active tension relaxation. **h,I**, Simulation results (**h**) that consider the myosin contribution to thin filament activation replicate the prolongation of the calcium transient decay observed experimentally (**i**). **j,k**, The combination of larger myosin availability and the myosin-based contribution to thin filament activation leads to a prolongation in the relaxation of simulated active tension (**j**) similar to experimental data (**k**).

Simulation results demonstrated a mechanistic link between myosin availability and hypercontractility in *MYH7*^R403Q/+^ (Fig. 2a-e), but not slowed cellular relaxation. We tested if increased myosin DRX availability could directly influence thin filament activation^17^ (Fig. 2f and Supplementary Fig. 3), and explain prolonged cellular relaxation. These simulations showed that myosin-based activation of the thin filament triggered slowed relaxation (Fig. 2g, darker colours). Thin filament activation by increased DRX myosin prolonged calcium transient (Fig. 2h,i) and active tension (Fig. 2j,k) decay in our in-silico models, as subsequently validated in *MYH7*^R403Q/+^ hiPSC-CMs (Fig. 2i,k).

### TNNT2^R92Q/+^ drives HCM by altering tropomyosin positioning and increasing calcium sensitivity

Altered tropomyosin positioning^18^ and increased calcium sensitisation of the thin filaments^19,20^ are intrinsic biophysical defects in *TNNT2*^R92Q/+^. We wanted to establish if both mechanisms are necessary to observe the cellular phenotype. When both factors are considered in our in-silico *TNNT2*^R92Q/+^ cardiomyocytes, we observe an increased tension amplitude and prolonged tension decay, consistent with increased amplitude (Fig. 3a,b) and prolonged relaxation (Fig. 3c,d) of sarcomere shortening in *TNNT2*^R92Q/+^ hiPSC-CMs. Simulated *TNNT2*^R92Q/+^ cardiomyocytes also replicated the prolonged calcium transient decay observed in *TNNT2*^R92Q/+^ hiPSC-CMs (Fig. 3e,f).

**Fig. 3.**
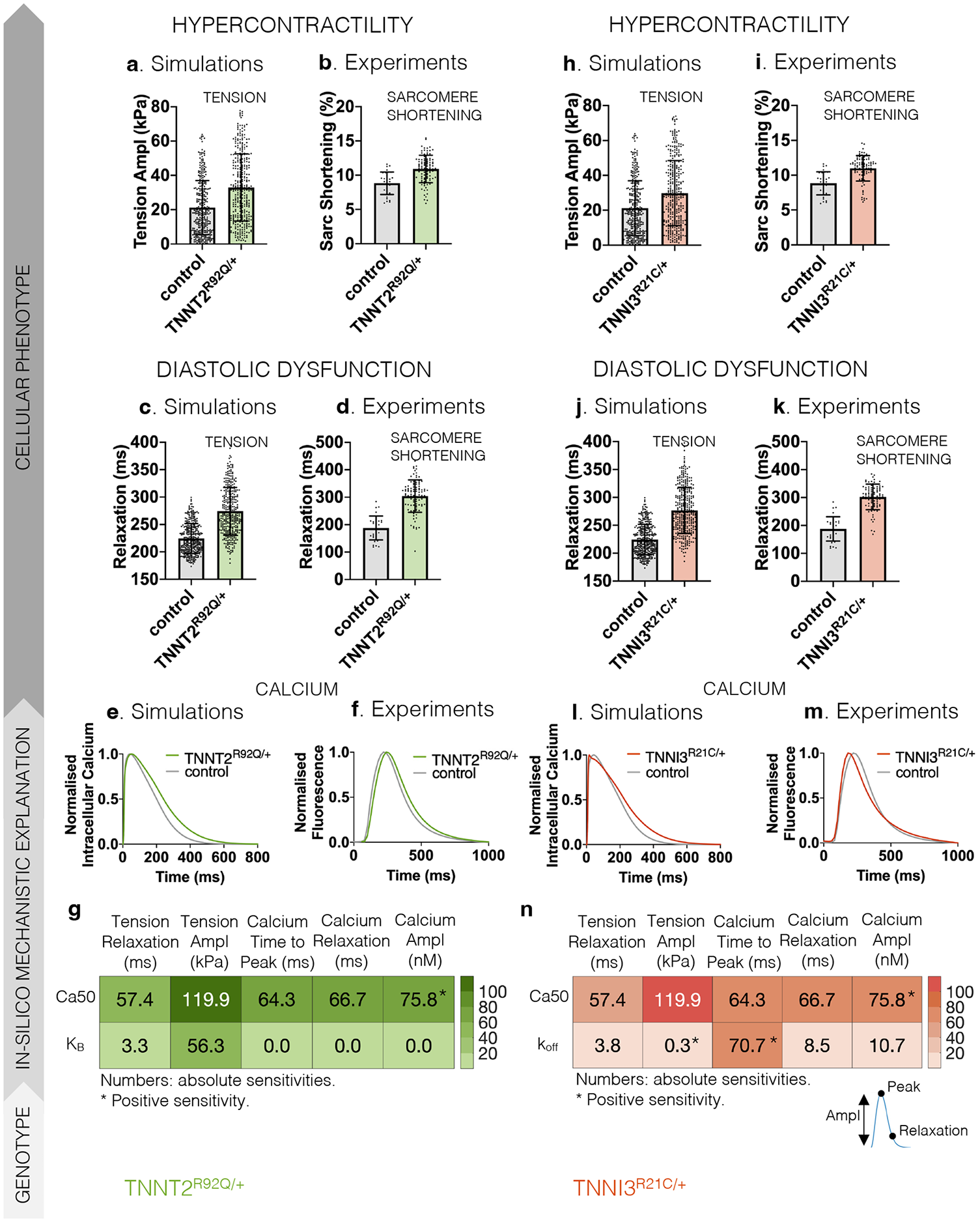
Mechanistic explanation of the TNNT2^R92Q/+^ and TNNI3^R21C/+^ cellular phenotypes of hypercontractility and diastolic dysfunction through human-based modelling and simulation informed by biophysical evidence. **a-d**, Simulated TNNT2^R92Q/+^ cardiomyocytes, which have altered tropomyosin positioning (K_B_) and increased calcium sensitivity (Ca50), develop higher tension amplitude (Tension Ampl, **a**) with prolonged relaxation time (Relaxation, **c**) compared to control, in line with experimental hiPSC-CM data (**b,d**). **e,f**, Simulation results (**e**) considering altered tropomyosin positioning and increased calcium sensitivity replicate the prolongation of the calcium transient decay observed experimentally (**f**). **g**, Absolute sensitivities of the active tension (Tension) and calcium transient (Calcium) biomarkers to changes in the model parameters that describe calcium sensitivity (Ca50) and tropomyosin positioning (K_B_). Biomarkers considered: Tension Relaxation denotes the time from tension peak to 90% decay, Tension Ampl denotes the difference between peak tension and baseline tension, Calcium Time to Peak denotes the time to reach peak calcium, Calcium Relaxation denotes the time from calcium peak to 90% decay, and Calcium Ampl denotes the difference between peak calcium and baseline calcium. **h-k**, Simulated TNNI3^R21C/+^ cardiomyocytes, which have altered calcium sensitivity (Ca50) and dissociation rate from troponin (k_off_), develop higher tension amplitude (**h**) with prolonged relaxation time (**j**) compared to control, in line with experimental hiPSC-CM data (**i,k**). **l,m**, Simulation results (**l**) considering altered calcium sensitivity and dissociation rate from troponin replicate the accelerated calcium rise and decelerated decay observed experimentally (**m**). **n**, Absolute sensitivities of the active tension (Tension) and calcium transient (Calcium) biomarkers to changes in the model parameters that describe calcium sensitivity (Ca50) and calcium dissociation rate from troponin (k_off_).

We used sensitivity analysis to establish the individual contribution of the two biophysical mechanisms to the HCM phenotype. To describe tropomyosin positioning, we modified the model parameter for the reverse rate constant of the equilibrium constant (K_B_) between blocked and closed tropomyosin states^18^. To regulate myofilament calcium sensitivity, we varied the parameter Ca50 describing the calcium concentration at half maximal thin filament activation. Ca50 had a larger impact on tension and calcium transient biomarkers compared to K_B_ (Fig. 3g). Both Ca50 and K_B_ contributed to changes in contractility (Fig. 3g, Tension Ampl). However, changes in tension and calcium relaxation were predominantly driven by changes in Ca50 (Fig. 3g, Tension Relaxation and Calcium Relaxation).

### TNNI3^R21C/+^ drives HCM by altering calcium binding and dissociation from troponin

*TNNI3*^R21C/+^ hiPSC-CMs exhibit an accelerated calcium rise and slowed calcium decay, hypercontractility, and prolonged relaxation^11^. We investigated whether the cellular *TNNI3*^R21C/+^ phenotype can be explained by the altered binding of calcium to troponin observed in^21^ driven by altered interactions between the mutated troponin I and troponin C. We considered altered Ca50 at the thin filament, as well as slower calcium dissociation (k_off_) from troponin. These variables explained increased active tension amplitude and prolonged tension decay time in-silico, as respectively validated by increased amplitude (Fig. 3h,i) and prolonged relaxation (Fig. 3j,k) of sarcomere shortening in *TNNI3*^R21C/+^ hiPSC-CMs. Calcium transients of simulated *TNNI3*^R21C/+^ cardiomyocytes showed accelerated calcium rise and slowed decay, as observed in *TNNI3*^R21C/+^ hiPSC-CMs (Fig. 3l,m).

Sensitivity analysis explained the relative contributions of calcium sensitivity Ca50 and calcium dissociation k_off_ to the *TNNI3*^R21C/+^ phenotype. Ca50 has a larger impact on both active tension and calcium transient biomarkers than alterations of k_off_ (Fig. 3n). Alterations in k_off_ were key to replicating the *TNNI3*^R21C/+^ phenotype of accelerated calcium rise (Fig. 3n, Calcium Time to Peak).

### Human in-silico trials explain the cellular mechanisms of Mavacamten’s efficacy in thin and thick filament HCM

We used our in-silico findings to investigate pharmacological efficacy of targeted therapeutics in HCM. We performed in-silico trials to simulate the effect of Mavacamten on *MYH7*^R403Q/+^, *TNNT2*^R92Q/+^, and *TNNI3*^R21C/+^ variants. Mavacamten has been shown to improve cellular HCM phenotypes in *MYBPC3* and *MYH7* gene variants through stabilisation of myosin SRX^13,16^. Mavacamten rescued both hypercontractility and impaired relaxation in *MYH7*^R403Q/+^ simulations. It dose-dependently reduced active tension of simulated *MYH7*^R403Q/+^ cardiomyocytes (Fig. 4a) by directly reducing myosin DRX, which deactivated knock-on thin filament activation in *MYH7*^R403Q/+^, showing correction of slowed relaxation (Fig. 4b). However, Mavacamten only partially corrected the contractile and prolonged relaxation phenotypes of *TNNT2*^R92Q/+^ (Fig. 4c,d) and *TNNI3*^R21C/+^ (Fig. 4e,f) cells. In our simulations Mavacamten can restore contractile function, irrespective of the mechanism that drives hypercontractility. However, correction of abnormal relaxation was only evident in models with overactive myosin, as found in *MYH7*^R403Q/+^. These in-silico predictions were borne out by our hiPSC-CM studies^11-13^ (Fig. 4g,h), where Mavacamten normalised cellular hypercontractility in all variants. However, it only corrected impaired relaxation at 3μM in *TNNI3*^R21C/+^ and not at all for the *TNNT2*^R92Q/+^ variant, compared to 0.5μM Mavacamten in *MYH7*^*403Q/+*^ hiPSC-CMs. Depression of contractility at 3μM Mavacamten was marked in the *TNNI3*^R21C/+^ variant, suggesting this dosage may fall outside the therapeutic window.

**Fig. 4.**
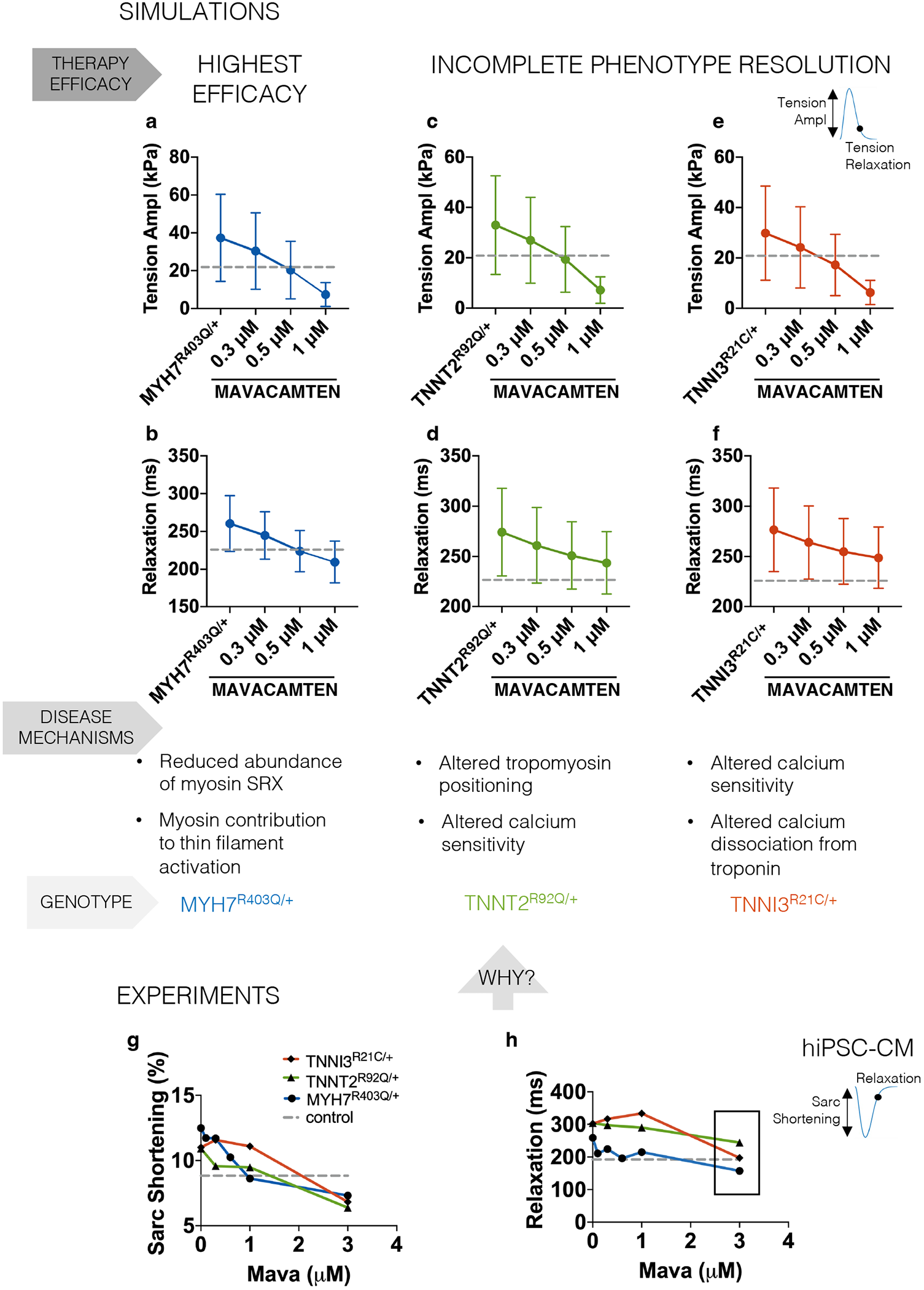
In-silico trials of Mavacamten on simulated MYH7^R403Q/+^, TNNT2^R92Q/+^, and TNNI3^R21C/+^ cardiomyocytes. **a,b**, Dose-dependent effect of Mavacamten on the active tension amplitude and relaxation time of simulated MYH7^R403Q/+^ cardiomyocytes, which are characterised by larger myosin availability due to a lower abundance of myosin SRX that contributes to thin filament activation. **c,d**, Dose-dependent effect of Mavacamten on the active tension amplitude and relaxation time of simulated TNNT2^R92Q/+^ cardiomyocytes, which are characterised by altered tropomyosin positioning and calcium sensitivity. **e,f**, Dose-dependent effect of Mavacamten on the active tension amplitude and relaxation time of simulated TNNI3^R21C/+^ cardiomyocytes, which are characterised by altered calcium sensitivity and dissociation rate from troponin. Simulation results recapitulated the experimentally observed response to Mavacamten of hiPSC-CMs expressing the MYH7^R403Q/+^, TNNT2^R92Q/+^, and TNNI3^R21C/+^ variants (**g,h**), and provide an explanation based on mutation-specific disease mechanisms.

### Human whole-ventricular electromechanical simulations of genotype-specific clinical phenotypes

To bridge from bench to bedside, and test Mavacamten in a whole-organ in-silico clinical trial, we simulated the clinical phenotypes of two virtual genotype-positive phenotype-negative (i.e. before left ventricular (LV) hypertrophy) HCM subjects carrying the *MYH7*^R403Q/+^ and *TNNT2*^R92Q/+^ variants (Fig. 5a,b). We do this to separate and understand the disease pathomechanism in isolation from the downstream organ-level adaptations and disease progression pathways. Therefore, we used the same ventricular anatomy (chamber size and wall thickness) for the two virtual subjects and only varied cellular properties. Human whole-ventricular electromechanical simulations showed that the *MYH7*^R403Q/+^ variant causes increased ventricular contractility, in good agreement with *MYH7*^R403Q/+^ cells (Fig. 5c). This is evident in the pressure-volume relation, driven by a smaller LV end-systolic volume in *MYH7*^R403Q/+^ of 65ml versus 77ml in healthy ventricles. LV end-diastolic volume was also reduced to 154ml in *MYH7*^R403Q/+^ versus 165ml in healthy ventricles. This signifies early-stage diastolic impairment. Overall, the LV ejection fraction (LVEF) increases in *MYH7*^R403Q/+^ compared to healthy control (58% vs 53%).

**Fig. 5.**
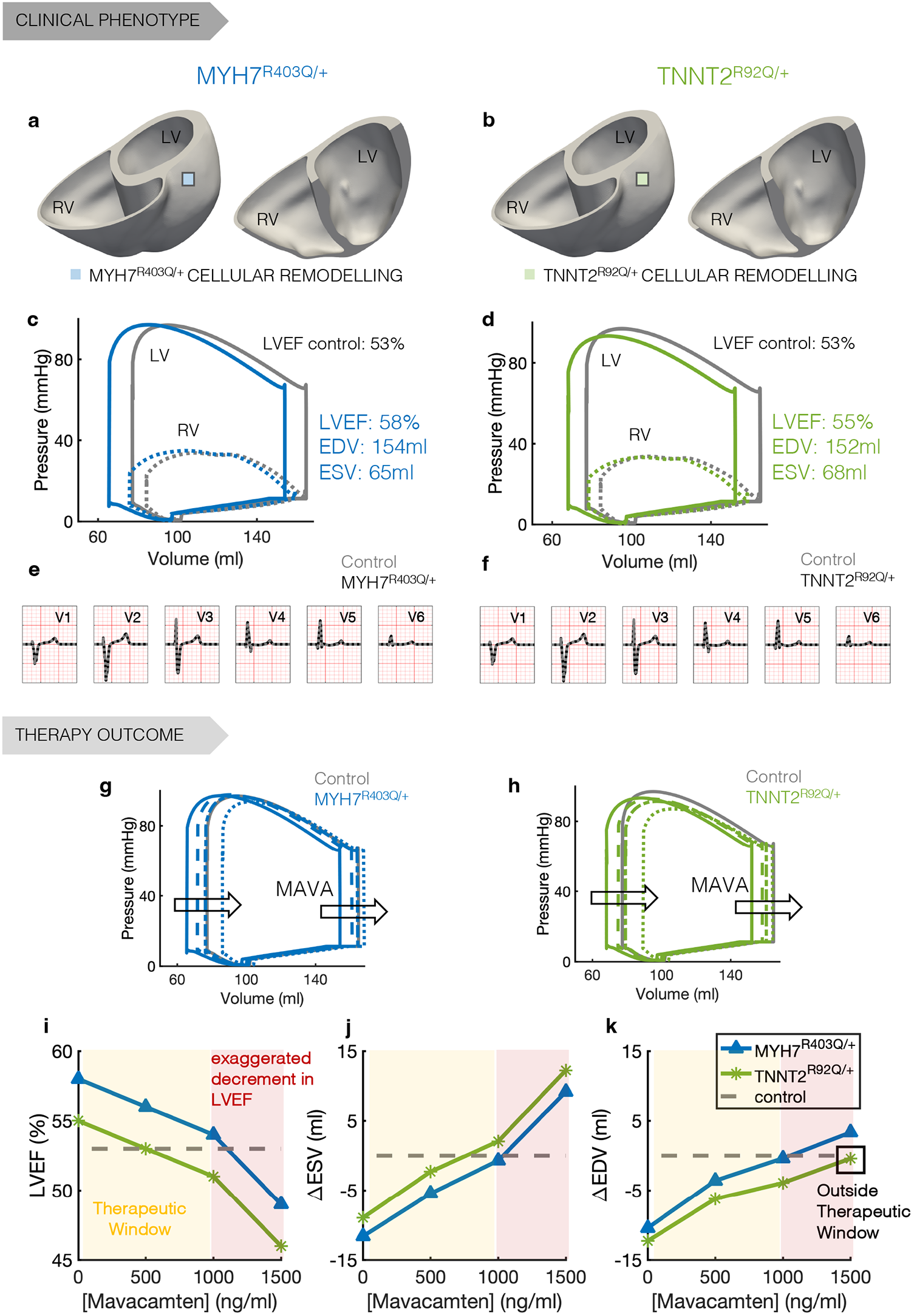
Human in-silico clinical trials of Mavacamten on genotype-positive phenotype-negative HCM virtual subjects. **a,b**, Magnetic resonance derived anatomical model for the genotype-positive phenotype-negative HCM virtual subjects. Mutation-specific cellular remodelling is homogeneously distributed throughout the ventricles. **c,d**, Pressure-volume loops of the left (solid lines) and right (dotted lines) ventricles of the virtual HCM subjects expressing the MYH7^R403Q/+^ (**c**, blue) and TNNT2^R92Q/+^ (**d**, green) genetic variants compared to control (grey). **e,f**, Comparison of the 12 lead ECGs of the HCM subjects expressing the MYH7^R403Q/+^ (**e**) and TNNT2^R92Q/+^ (**f**) genetic variants compared to control (grey). **g,h**, Dose-dependent effect of Mavacamten (500, 1000, and 1500ng/ml) on the pressure-volume loop of the left ventricle of the HCM subjects expressing the MYH7^R403Q/+^ (**g**) TNNT2^R92Q/+^ (**h**) genetic variants. **i-k**, Comparison of the dose-dependent effects of Mavacamten on the LVEF (**i**), end-systolic volume (**j**), and end-diastolic volume (**k**) of the HCM subjects expressing the MYH7^R403Q/+^ (blue) and TNNT2^R92Q/+^ (green) genetic variants. The yellow area represents the therapeutic window of the drug whereas the red area identifies drug concentrations that lead to a LVEF below 50%.

The *TNNT2*^R92Q/+^ variant virtual ventricles also displayed hypercontractility due to a smaller LV end-systolic volume (68ml vs 77ml of a healthy subject, Fig. 5d). LV end-diastolic volume was reduced to 152ml in the *TNNT2*^R92Q/+^ versus 165ml in healthy ventricles. The reduction in end-systolic volume was more pronounced in *MYH7*^R403Q/+^ than in *TNNT2*^R92Q/+^, while the reduction of end-diastolic volume was more pronounced in *TNNT2*^R92Q/+^ than in *MYH7*^R403Q/+^. Overall, *MYH7*^R403Q/+^ ventricles present a higher LVEF of 58% compared to the 55% of *TNNT2*^R92Q/+^ (Fig. 5c,d). Our model shows a stronger hypercontractility phenotype in *MYH7*^R403Q/+^, while *TNNT2*^R92Q/+^ ventricles exhibit a more severe diastolic dysfunction. In both cases, despite phenotypic ventricular changes in mechanical function typical of early genotype-positive phenotype-negative HCM^22^, simulated 12-lead ECGs did not show signs of abnormalities in the virtual patients (Fig. 5e,f) as occurs in the majority of preclinical subjects^23^.

### Mavacamten shows incomplete rescue of thin filament HCM in phenotype-negative virtual ventricles

Our cellular results demonstrate Mavacamten’s efficacy on variants that alter myosin SRX but suggest that variants not altering myosin availability (*TNNI3, TNNT2*) may receive a more modest cellular therapeutic benefit. We tested this hypothesis in a whole-organ in-silico clinical trial. We tested the effect of clinically-relevant plasma concentrations of Mavacamten at 500, 1000, and 1500ng/ml on the electromechanical function of the *MYH7*^R403Q/+^ and *TNNT2*^R92Q/+^ virtual ventricles. We observed a dose-dependent correction of hemodynamics in *MYH7*^R403Q/+^ and *TNNT2*^R92Q/+^ carriers (Fig. 5g,h). This translated into a dose-dependent reduction of LVEF, which remained above the safety threshold (LVEF>50%) up to 1000ng/ml Mavacamten (Fig. 5i). Concentrations above 1000ng/ml caused an exaggerated reduction in LVEF that may not be clinically desirable^24^. We observed a marked reduction of LVEF from 54% to 49% in *MYH7*^R403Q/+^ and from 51% to 46% in *TNNT2*^R92Q/+^ ventricles, in the 1000-1500ng/ml Mavacamten range. Our results support the safety of Mavacamten when administered in the therapeutic range of 350-700ng/ml in line with clinical findings^10^. Importantly, simulation results highlight differences in therapeutic efficacy. The abnormal end-systolic volume is dose-dependently corrected within the therapeutic window in both virtual variant carriers (Fig. 5j), but Mavacamten only restored the end-diastolic volume within the therapeutic window of *MYH7*^R403Q/+^ but not *TNNT2*^R92Q/+^ ventricles (Fig. 5k), in agreement with our cellular results. Altogether, our organ simulations underline the efficacy of Mavacamten for genetic variants that alter myosin conformations, such as *MYH7*^R403Q/+^, but further suggest an unmet need in correction of cell to organ abnormalities caused by thin filament HCM, which we investigate below.

### Human in-silico trials identify pathomechanism-targeted thin filament HCM therapeutics

Given the incomplete correction of HCM cellular and organ level phenotypes by Mavacamten in thin filament variants, we trialled additional pharmacological targets to establish the efficacy of directly targeting primary pathomechanisms in this subset of HCM carriers.

We tested on-market L-type calcium and late sodium current blockers^25^, as well as upregulation of sarcoplasmic endoplasmic reticular calcium ATPase (SERCA), as SERCA is reduced in human HCM samples^26-28^. The effect of late sodium current block on relaxation in *TNNT2*^R92Q/+^ and *TNNI3*^R21C/+^ cardiomyocytes was minimal, with partial correction of hypercontractility (Supplementary Fig. 4a,b). A similar trend was observed for L-type calcium current block (Supplementary Fig. 4c,d), although severe negative inotropic effects were present. SERCA upregulation restored tension relaxation, despite an undesirable further positive inotropic effect (Supplementary Fig. 4e,f). Only dual application of SERCA upregulation and 0.5μM Mavacamten achieved complete phenotype resolution (Supplementary Fig. 4e,f, triangles).

Clinical upregulation approaches are often complicated by delivery strategies, which would be further complicated by trialling a non-targeted polytherapeutic/polypharmacy approach by the addition of Mavacamten. We hypothesised that a single-target therapeutic could provide a more efficacious therapeutic strategy than a multi-target one. Therefore, we generated a framework to define a novel thin filament-specific drug. We took advantage of the finding that diastolic dysfunction in *TNNT2*^R92Q/+^ and *TNNI3*^R21C/+^ variants are both due to increased myofilament calcium sensitivity. We tested an idealised pharmacological intervention specifically acting as a calcium desensitiser at the thin filament (Fig. 6a). This strategy dose-dependently normalised both contraction and relaxation in thin filament HCM (Fig. 6b,c), leading to phenotype resolution in simulated *TNNT2*^R92Q/+^ cardiomyocytes at a 50% calcium sensitivity reduction (Fig. 6d,e). A 50% decrease in calcium sensitivity at the organ level also restored both systolic and diastolic ventricular dysfunction without deleterious effects on LVEF (Fig. 7f,g). Both end-diastolic (Fig. 6h) and end-systolic (Fig. 6i) volumes were corrected within therapeutic window.

**Fig. 6.**
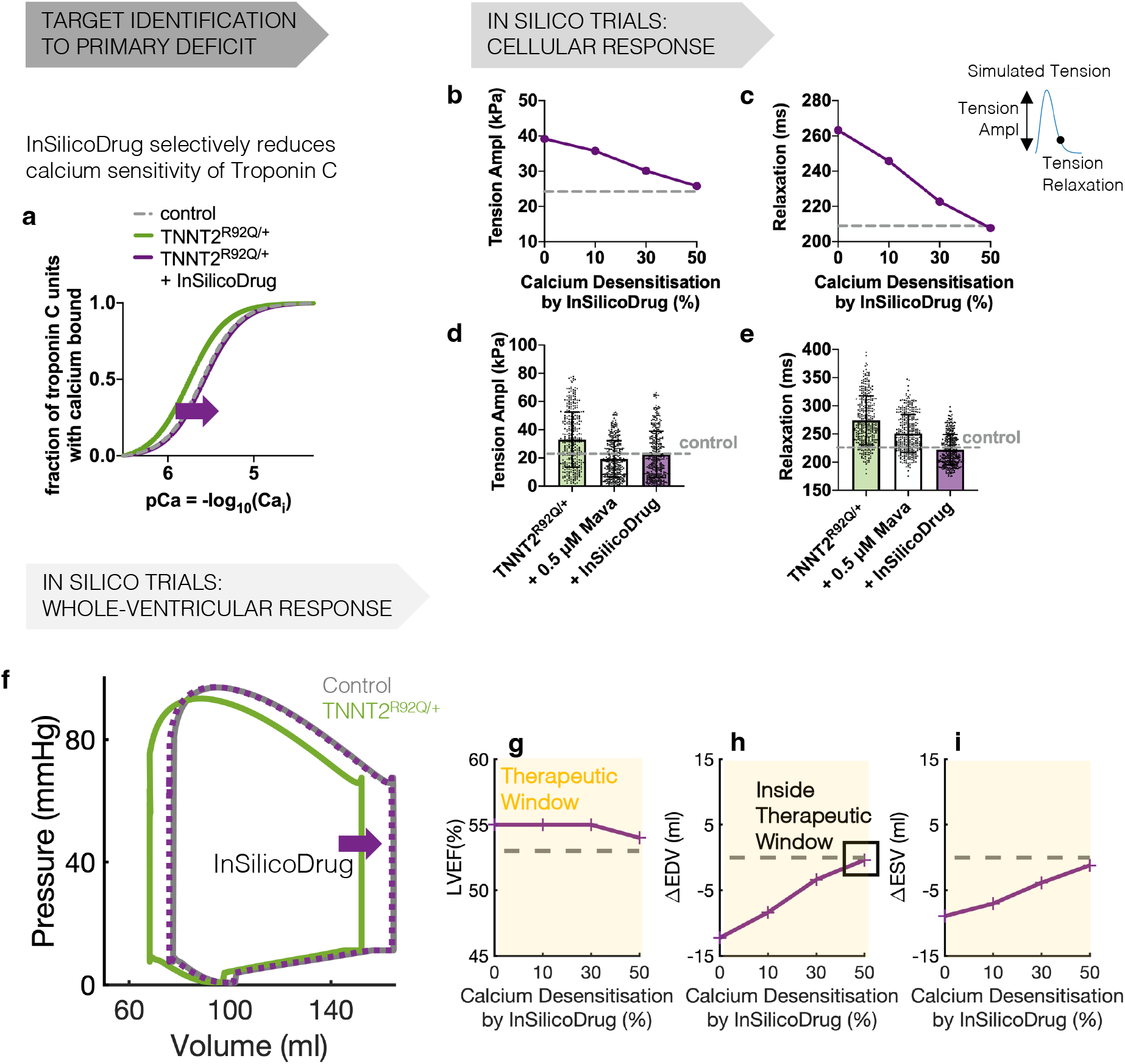
In-silico identification of sarcomere druggable targets for the resolution of the TNNT2^R92Q/+^ phenotype when Mavacamten’s action only is suboptimal. **a**, Based on identified pathophysiology, the in-silico designed drug should selectively reduce calcium sensitivity of troponin C and shift the steady state relationship between calcium bound to troponin and free calcium to the right. **b,c**, Effect of the in-silico designed drug on tension amplitude (**b**) and relaxation (**c**) of a simulated TNNT2^R92Q/+^ cardiomyocyte. **d,e**, Comparison of the effects of 0.5μM Mavacamten and the in-silico designed drug (50% calcium desensitisation of the thin filament) on the tension amplitude (**d**) and relaxation (**e**) of a population of simulated TNNT2^R92Q/+^ cardiomyocytes. **f**, TNNT2^R92Q/+^ pressure-volume loop normalisation by the in-silico designed drug. **g-i**, Dose-dependent effect of the in-silico designed drug on the LVEF (**g**), end-diastolic (**h**), and end-systolic (**i**) volume of the left ventricle of the virtual TNNT2^R92Q/+^ carrier. Yellow areas identify therapeutic windows. The in-silico identified drug restores all phenotype components within therapeutic window.

## Discussion

Within the field of genetic cardiomyopathy and more widely in cardiovascular research there have been many techniques developed to accelerate variant classification and therapeutic screening^13,29,30^, spanning biochemical investigations, rapid variant classification platforms, and hiPSC-CM-based systems. But with ever-growing genomic datasets, biological/biochemical phenomics is still a bottleneck in translation from bench to bedside. Herein we describe a synergistic hybrid biological and in-silico human system that accelerates hypothesis generation and testing in variant classification, pathomechanism discovery, and therapeutic targeting. Our human modelling and simulation approach bridges from disease-causing mutations to clinical disease manifestation. We address hiPSC-CMs immaturity through the modelling of human adult virtual cardiomyocytes, and physiological variability through populations of models. We scale this analysis to genotype-specific clinical phenotypes which are obtained through whole-ventricular simulations. We used human-based digital twins and mechanistic simulations to understand pathomechanisms in HCM and identify druggable targets. Altogether, this generates digital evidence of incomplete rescue of thin filament HCM phenotypes by Mavacamten but shows the promise of combination therapies or novel targeted therapeutics.

hiPSC-CMs provide a human cellular system to validate these findings^31,32^, as they capture patient-specific genotype-phenotype relationships and pharmacological responses^33^. hiPSC-CM experiments show that both thick (*MYH7*^R403Q/+^) and thin (*TNNT2*^R92Q/+^, *TNNI3*^R21C/+^) filament HCM variants cause cellular hypercontractility and diastolic dysfunction^11-13^, consistent with clinical findings in HCM^10,22,34^. Our simulations show that SRX destabilisation is central to hypercontractility in thick filament HCM variants in *MYH7*^13,29,35^, but cannot explain impaired relaxation. We found that, secondarily to myosin activation, knock-on thin filament activation prolongs calcium binding to troponin^17^, slowing cellular relaxation in the *MYH7*^R403Q/+^ phenotype. When applying these mechanisms to whole-ventricular simulations, the *MYH7*^R403Q/+^ ventricles presented lower end-diastolic and end-systolic volumes and increased LVEF, consistent with clinical findings^22,34^.

We found that the thin filament HCM variants studied here have changes in thin filament activation driven by altered calcium sensitivity^19^. *TNNT2*^R92Q/+^ sensitises myofilament calcium binding by alterations to cooperative activation^19^, where its initial biophysical insult could reside in tropomyosin positioning^18^. Our in-silico models confirm this and show indirect calcium sensitisation of the myofilament as a key determinant of diastolic dysfunction and calcium transient abnormalities in *TNNT2*^R92Q/+19,20,36^. This is confirmed by whole-ventricular simulations, where the increased myofilament calcium sensitivity drives a greater diastolic insufficiency in *TNNT2*^R92Q/+^ than in *MYH7*^R403Q/+^ ventricles.

*TNNI3*^R21C/+^ hiPSC-CMs exhibit faster calcium transient rise and prolonged transient decay, alongside hypercontractility and impaired relaxation^11^. We show that the abnormal calcium transient phenotype is caused by increased calcium binding and slowed dissociation from troponin, leading to greater crossbridge recruitment, hypercontraction, and prolonged relaxation^21^.

Mavacamten, an allosteric myosin ATPase inhibitor, was recently approved by the FDA for the treatment of symptomatic obstructive HCM. Developed to target the core molecular mechanisms that cause HCM^9^, Mavacamten reduces ATPase activity and increases myosin SRX formation across a variety of experimental models9^,13,16,35,37-41^. Building on positive preclinical results, Mavacamten was advanced to clinical testing^10,24,42^, but the exact mechanisms through which it restores contractile function, especially across HCM genotypes, are not fully understood. To investigate this, we extended our cellular human model to integrate modulation of myosin availability.

Application of Mavacamten to our *MYH7*^R403Q/+^ thick filament HCM models predicted a clear resolution of cellular HCM phenotypes, in agreement with our *MYH7*^R403Q/+^ hiPSC-CM data. Thin filament HCM was not fully rescued by Mavacamten, confirmed using *TNNT2*^R92Q/+^ and *TNNI3*^R21C/+^ hiPSC-CMs^11-13^. Mavacamten was able to dose-dependently suppress cellular contractility in-silico irrespective of genotype, but its efficacy in restoring diastolic function was diminished in thin filament variants. We provide digital evidence that this is because Mavacamten does not directly target the HCM causative mechanisms in these variants. Mavacamten’s administration in ventricular simulations of thick filament genotype-positive phenotype-negative HCM reverses early phenotypic abnormalities of hyperdynamic ventricular contraction and impaired ventricular filling, with modest LVEF decrease. This suggests that Mavacamten can potentially be used as a treatment to prevent disease progression, in agreement with preclinical testing^9^. Our simulations predicted a tolerable decrease in LVEF at 350-700ng/ml Mavacamten with detrimental reductions in LVEF at 1000ng/ml, as previously reported^10,24^. Simulations show that hyperdynamic ventricular contraction is corrected by Mavacamten in thin filament variants. However, Mavacamten only showed correction of impaired ventricular filling in *MYH7*^R403Q/+^ and not *TNNT*^R92Q/+^. In agreement with our cellular results, whole-organ diastolic insufficiency caused by thin filament HCM cannot be corrected by Mavacamten without detrimental suppression of LVEF. Our study highlights the importance of future investigations into Mavacamten’s utility in specific HCM genotype populations.

We define the importance of a pharmacogenetic approach that defines and targets the incident mechanism of pathophysiology in thin filament HCM, which is an increased myofilament calcium sensitivity. We show this is feasible by selective calcium desensitisation, which restores cellular function and normalises whole-organ contractility without detrimental suppression of LVEF in *TNNT2*^R92Q/+^. This highlights the potential of hybrid human systems as powerful tools to accelerate the development and administration of rational therapies in a pathomechanism-specific manner.

## Methods

### Human in-vitro single cell model

Heterozygous pathogenic missense variants *TNNI3*^R21C/+^, *TNNT2*^R92Q/+^ and *MYH7*^R403Q/+^ that cause HCM were introduced using CRISPR/Cas9 technology as previously described in an hiPSC cell line harbouring GFP-labelled titin^43,44^. Monolayer differentiation of control and WT cell lines was performed via Wnt pathway modulation with small molecule inhibitors. Cells were induced to the mesodermal layer with 12μM CHIR99021 in RPMI1640 / B27 minus insulin for 24 hours. This was considered to be day 0 of differentiation. At the end of 24 hours, CHIR99021 was diluted to 6μM with fresh RPMI1640 / B27 minus insulin for a further 24 hours. On day 2, media were replaced with RPMI1640 / B27 minus insulin. On day 3, cells were induced to cardiac lineage specification with 5μM IWP2 in RPMI1640 / B27 minus insulin for 48 hours. Cells were then cultured in the absence of insulin to day 10. Spontaneous contraction was observed on day 9-11 of differentiation. Cells were subjected to two 48-hour rounds of metabolic selection starting on day 10, with glucose-free RPMI / B27 plus insulin. Cells were then cultured to day 24 in RPMI1640 / B27 plus insulin. On day 24, cells were passaged onto glass-bottom dishes and cultured to day 28 in RPMI1640 / B27 plus insulin. On day 28, cells designated for calcium phenotyping were induced to express the red genetically encoded calcium indicator for optogenetics, RGECO1, via adenoviral transduction at MOI 40 for 24 hours. The next day, media were changed back to RPMI1640 / B27 plus insulin. Plates designated for contractility measurement were cultured to day 30 in to RPMI1640 / B27 plus insulin.

All measurements were prepared in triplicate technical replicates and triplicate biological replicates. Differentiations of separate cell passages were considered distinct biological replicates. Cells designated for contractility measurement were incubated for ten minutes at 37°C / 5% CO_2_ with the myosin inhibitor Mavacamten at 0.3, 1 and 3μM prior to imaging in Tyrodes-HEPES buffer at 37°C / 5% CO_2_ and imaged directly. For calcium measurement, mutant and WT cells were equilibrated with Tyrode’s-HEPES buffer for ten minutes at 37°C / 5% CO_2_ and then imaged. Controls were provided by triplicate plates of equivalent concentrations of dimethylsulfoxide (DMSO) and triplicate plates incubated in the absence of any treatment to capture the mutant phenotype.

hiPSC-CMs from each group were imaged on an Olympus IX81 inverted microscope (Olympus, Japan) with a C-9100-13 EMCCD camera (Hamamatsu, Japan), under electrical stimulation of 20V / 1Hz and at 37°C. Videos were acquired at 50 fps, 560/25 nm excitation, 620/60 nm emission with a 565 nm dichroic mirror for calcium imaging and 50fps, 485/20-25nm excitation, 525/50nm emission with a 495nm dichroic mirror for contractility imaging. Contractility measurement for *MYH7*^R403Q/+^ was performed at 30fps. A minimum of 30 cells were recorded per plate. Data was extracted with CalTrack^11^ and SarcTrack^45^ software.

### Human in-silico single cell model

We extended our coupled human electromechanical adult cardiomyocyte model^14,15,46^ to allow interrogation of how disease- and drug-induced changes in the physiological regulation of myosin function alter ventricular contractility. We hypothesised that changes in myosin SRX could be phenomenologically simulated with the coupled model through a modulation of actin-myosin crossbridge availability. This was computationally implemented as an explicit dependency of crossbridge formation on the proportion of myosin in the SRX state, which are inhibited and unable to bind actin, with respect to the rest of myosin heads that are available to drive contraction, which broadly exist in the myosin disordered relaxed state DRX. We introduced a new parameter 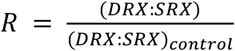 into the cellular model to account for deviations from the control DRX:SRX ratio. We calibrated the modified cellular model to achieve changes in contractile function by SRX:DRX modulation that are consistent with experimental data. Details of the cellular model construction and calibration are reported in the supplement.

### Populations of in-silico human adult cardiomyocytes with different genetic background

From the extended control single cell model, a control population of healthy cardiomyocytes was built to consider physiological electromechanical variability^47^. An initial population of 2,000 electromechanical models was created by sampling the fast and late sodium, transient outward potassium, rapid and slow delayed rectifier potassium, inward rectifier potassium, sodium-calcium exchanger, sodium-potassium pump, and the L-type calcium conductances, the sarcoplasmic reticulum calcium release flux, the calcium uptake via SERCA, calcium sensitivity of SERCA, intracellular sodium affinity of the sodium-potassium pump, calcium sensitivity of myofilament, and crossbridge cycling rates. Parameters were varied in the range [50-200]% of their baseline value with the Latin Hypercube Sampling technique. The population was calibrated based on action potential and calcium transient data as in^48^ and on active tension data (time to 50 and 95% transient decay). We considered experimental action potential, calcium transient, and active tension recordings data as in^15^.

The control population was then used to construct three populations with different genetic background, representative of the three HCM causing variants considered in this study. To build the *MYH7*^R403Q/+^ population, we introduced a remodelling in the myosin ratio 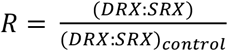 of 1.3, as measured experimentally in hiPSC-CMs^13^. To investigate the possible determinants of impaired cellular relaxation in *MYH7*^R403Q/+^, we compared simulation results obtained with this population with results from a different population which additionally considered a myosin-based contribution to thin filament activation (as described in the supplement). The *TNNT2*^R92Q/+^ population was constructed based on the reported early-stage changes in thin filament protein function due to the mutation, and the parameters of this were calibrated using the results of a sensitivity analysis. Mutation-induced early changes in thin filament protein function included altered tropomyosin positioning on actin^18^ and an indirect calcium sensitisation of myofilaments^19,20^. The sensitivity analysis determined how much model parameters related to these two factors, namely K_B_ and Ca50, should change in order to obtain the expected hypercontractility and impaired relaxation observed experimentally. Further details are reported in the supplement. The *TNNI3*^R21C/+^ population was constructed following a similar approach, as outlined in the supplement. Here we considered a direct calcium sensitisation of myofilaments (Ca50) as well as altered dissociation rate from troponin (k_off_), based on^21^.

### Human-based in-silico drug trials

We used our generated cellular populations to run in-silico trials of pharmacological therapies and identify effective targets based on identified pathomechanisms for each genotype. We simulated the novel myosin modulator Mavacamten by using the relationship we established between Mavacamten concentrations and myosin availability when we calibrated our cellular model (as explained in the supplement). This provided a dose-response curve through which values for the model parameter 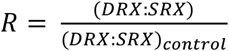 can be estimated from Mavacamten concentrations, enabling its testing in virtual trials. We simulated Mavacamten concentrations in the experimentally relevant range of 0.3-1.0μM^9,13,49^. We computed biomarkers of active contraction to determine the effect of the drug on the contractility and diastolic function of simulated *MYH7*^R403Q/+^, *TNNT2*^R92Q/+^, and *TNNI3*^R21C/+^ cardiomyocytes, and compared simulated results with experimental evidence^11-13^. Specifically, we considered the amplitude of the tension transient, and the relaxation time quantified as time to 90% transient decay of the simulated twitch tension.

In order to identify additional therapeutic targets that could aid in phenotype resolution for the troponin variants, we also tested the effect of additional pharmacological strategies. Specifically, we tested different levels of L-type calcium and late sodium currents block (20, 40, 60% block of current conductances) and different levels of SERCA upregulation (20, 50, 80% increase) in the presence and absence of 0.5μM Mavacamten. We computed biomarkers of active contraction to determine the effect of these simulated therapeutic strategies on the contractility and diastolic function of simulated *TNNT2*^R92Q/+^ and *TNNI3*^R21C/+^ cardiomyocytes. Finally, we designed in-silico a pharmacological intervention that, through a direct calcium desensitisation of the myofilaments, could rescue the identified mechanisms of disease caused by the *TNNT2*^R92Q/+^ and *TNNI3*^R21C/+^ genetic variants, thereby restoring the associated disease phenotype. We tested different percentages (10, 30, and 50%) of calcium desensitisation by the in-silico identified drug on the cellular and organ function under *TNNT2*^R92Q/+^. We computed cellular and organ level mechanical biomarkers and compared therapeutic efficacy against Mavacamten.

### Human whole-ventricular electromechanical simulations

We conducted in-silico clinical trials of Mavacamten through human biventricular electromechanical simulations to investigate implications on clinical phenotypes.

We used a recently published biventricular model that features a torso-biventricular anatomy from clinical magnetic resonance imaging^50^. The torso-biventricular mesh features an average element edge length of 220 μm. This allowed to compute pressure and volume transients and 12-lead electrocardiograms at clinically standardised lead locations as in^50^ and to extract clinical electrocardiograms biomarkers, end-diastolic and end-systolic volumes, and left and right ventricular ejection fractions.

Here we integrated our extended cellular electromechanical model, in control conditions and under the *MYH7*^R403Q/+^ and *TNNT2*^R92Q/+^ variants, to construct the digital twin of healthy and genotype-positive phenotype-negative HCM subjects. The scaling factor of cellular active tension T_scale_ in our biventricular model^50^ was calibrated to achieve 53% LVEF in control conditions, and set to 4. We conducted biventricular simulations as described in^50^ and studied how the changes in cellular contractile function driven by the *MYH7*^R403Q/+^- and *TNNT2*^R92Q/+^-specific remodelling affected whole-ventricular contractility.

To be able to conduct in-silico clinical trials of Mavacamten, we calibrated our cellular model of the drug with clinical data from healthy subjects as described in the supplement (Supplementary Fig. 5). Our calibrated model of Mavacamten allowed the estimation of the drug free plasma concentrations to be tested in-silico to simulate clinically administered drug doses. We used the calibrated model to test the dose-dependent effect of 500, 1000, and 1500ng/ml Mavacamten (based on clinical trials^10,24,42^) on whole-ventricular electromechanical function. Simulated clinical biomarkers were compared with clinical evidence^10,24,42^.

### Software and stimulation protocols

Cellular simulations were conducted as in^15^ using MatLab (Mathworks Inc. Natick, MA, USA) and the ordinary differential equation solver ode15s. In each simulation, we delivered a stimulus current of -53μA/μF with 1ms duration, and considered a fixed extension ratio (λ, sarcomere length over sarcomere length at rest) of 1. For each simulation, steady-state was reached at 1 Hz pacing before biomarkers were computed. Simulations of populations of models were conducted using the University of Oxford Advanced Research Computing (ARC) facility.

Whole ventricular simulations were conducted as in^50^, using the high-performance numerical software Alya for complex coupled multi-physics and multi-scale problems^51^ on the supercomputer *Piz Daint* of the Swiss National Supercomputing Centre. We simulated three beats of 800 ms cycle length with cellular models in steady-state conditions, and the pressure-volume loop and ECG convergence were assessed before computing clinically relevant endpoints.

## Acknowledgments

F. Margara was funded by the Personalised In-Silico Cardiology project (European Union’s Horizon 2020 research and innovation programme under the Marie Sklodowska-Curie grant agreement 764738) and by an Engineering and Physical Sciences Research Council Impact Acceleration Account Award EP/R511742/1. Support for this study was provided by British Heart Foundation (BHF) and Wellcome Trust (BHF Intermediate Basic Science Fellowship (FS/17/22/32644) to A. Bueno-Orovio; Sir Henry Dale Wellcome Fellowship (222567/Z/21/Z) and BHF Centre of Research Excellence Intermediate Transition Fellowship (RE/18/3/34214) to C.N. Toepfer; Wellcome Trust Senior Research Fellowship (214290/Z/18/Z) to B. Rodriguez). A. Bueno-Orovio and B. Rodriguez also acknowledge support from an NC3Rs (National Centre for the Replacement, Refinement and Reduction of Animals in Research) Infrastructure for Impact Award (NC/P001076/1) and the Oxford BHF Centre of Research Excellence (RE/13/1/30181). Y. Psaras was funded by the BHF CRE (RE/18/3/34214). Additional support was provided by the Sarnoff Foundation and BHF (RG/12/16/29939) for G.G. Repetti; the Fondation Leducq for J.G. Seidman and C.E. Seidman; the Engineering Research Centers Program of the National Science Foundation (NSF) for J.G. Seidman and C.E Seidman (NSF Cooperative Agreement number EEC-1647837); the National Institutes of Health for J.G. Seidman and C.E. Seidman (5R01HL080494 and 5R01HL084553); and the Howard Hughes Medical Institute for C.E. Seidman. We acknowledge PRACE for awarding access to the Fenix Infrastructure resources at the Swiss National Supercomputing Centre, Switzerland (PRACE-ICEI grants icp005 and icp013), which are partially funded from the European Union’s Horizon 2020 research and innovation programme through the ICEI project under the grant agreement No. 800858. The authors also acknowledge the use of the University of Oxford Advanced Research Computing (ARC) facility in carrying out this work (http://dx.doi.org/10.5281/zenodo.22558).

## Supplementary Information

Supplementary information document

## Data availability

Cellular model code and Alya executable and simulation input files required to replicate the simulated results of this study, and hiPSC-CM experimental datasets reported in this study will be made available upon request.

## Author contributions

F.M. developed the cellular in-silico model, ran cellular and biventricular simulations, analysed simulation results and experimental data. Z.J.W. and R.D. developed the biventricular framework. Y.P. M.S. A.G. and G.G.R. performed the experiments. A.B-O. C.N.T. B.R. C.S. J.S. supervised the study. F.M. A.B-O. C.N.T. and B.R. designed the study and wrote the manuscript. All authors discussed the results and commented on the manuscript.

