## Supplementary_Information for "Human iPSC-CMs and in-silico technologies define mechanisms and accelerate targeted pharmacogenetics in hypertrophic cardiomyopathy"

### Supplement

**In-silico modelling of drug- and disease-induced myosin availability changes in human cardiomyocytes**

We extended our coupled human electromechanical cardiomyocyte model[1-3] to allow interrogation of how disease- and drug-induced changes in the physiological regulation of myosin function alter ventricular contractility. Our electromechanical model combines the *Tomek, Rodriguez - following ORd* (ToR-ORd) model of human ventricular electrophysiology and excitation-contraction coupling[4], and the *Land* model of active contraction[5]. It provides a biophysically detailed description of all key ionic processes such as ionic currents, exchangers, and pumps, and the excitation-contraction coupling system, allowing representation of cellular depolarisation, repolarisation, and properties of calcium dynamics[4]. The coupled model also features thin filament troponin C and tropomyosin kinetics, and a three-state acto-myosin crossbridge model to describe the cyclical interactions between actin and myosin that propel cardiac contraction[5].

We hypothesised that changes in the physiological occupancy of the myosin energy conserving super relaxed state SRX[6] could be phenomenologically simulated with the coupled model through a modulation of acto-myosin crossbridge availability. This was computationally implemented as an explicit dependency of crossbridge formation on the proportion of myosin in the SRX state, which are inhibited and unable to bind actin, with respect to the rest of myosin heads that are available to drive contraction, which broadly exist in the myosin disordered relaxed state DRX. We will refer to this proportion as the DRX:SRX ratio. Specifically, we introduced a new parameter into the cellular model to account for deviations from the control DRX:SRX ratio (Supplementary Fig. 1):

$$R = \frac{(DRX{:}SRX)}{(DRX{:}SRX)_{control}}$$

In human myocardium the majority of myosins reside in the SRX state resulting in $(\text{DRX:SRX})_{\text{control}}$ ~ 30:70[7,8]. This ratio can be biased towards the DRX state (R > 1) due to HCM genetic variants, and Mavacamten can shift DRX:SRX ratios towards SRX (R < 1). Human-based experimental evidence suggests that myosin DRX can vary between 20 and 60[7,8] and thus the R parameter has been simulated in the range $\frac{(20:80)}{(60:40)} < R < \frac{(60:40)}{(20:80)}$ $(0.2 < R < 6)$. To replicate contractility changes by myosin modulation in-silico, the model parameter R modulates how many acto-myosin crossbridges enter in the weakly-bound crossbridge state W, as stated in the equation below:

$$\frac{dW}{dt} = k_{uw} * f_1(R) * U - k_{wu} * f_2(R) * W - k_{ws} * W - \gamma_{wu} * W$$

This ordinary differential equation describes the time evolution of weakly-bound crossbridges (W), where $k_{uw}$, $k_{wu}$, $k_{ws}$, and $\gamma_{wu}$ represent transition rates, and U is the population of unblocked myosin binding sites on actin, or the detached crossbridge state. We refer to the original publication of further details on the model implementation[5]. $f_1(R)$ and $f_2(R)$ are novel functions introduced into the model to tune the contractile output based on myosin SRX:DRX proportions. Several mathematical considerations (as reported below) led us to choose the functions $f_1(R)$ and $f_2(R)$ to be of the form $f_i(R) = R^n$.

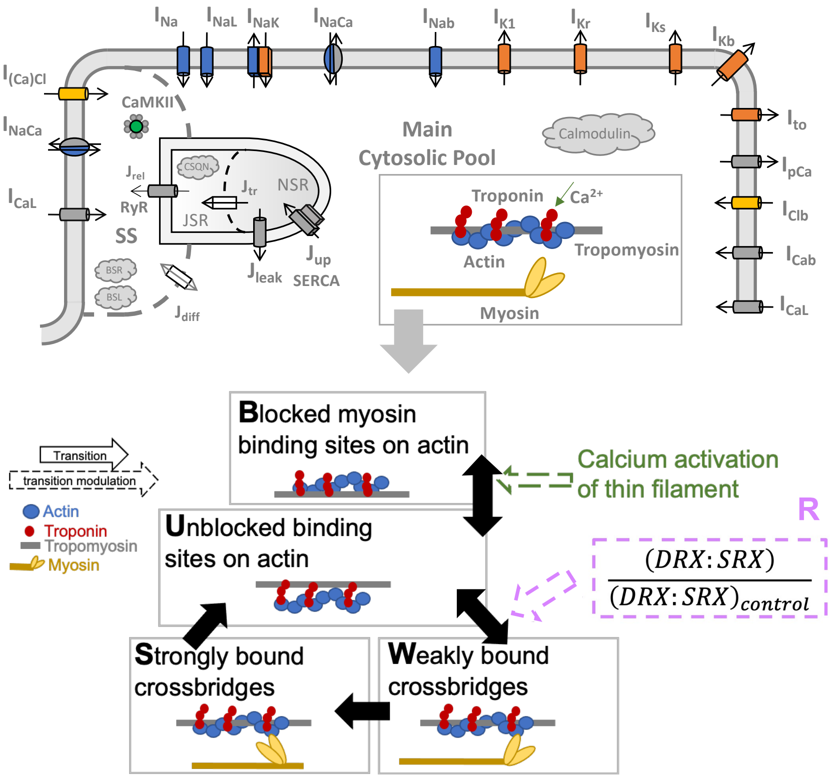


Supplementary Fig. 1. Schematic of the extended single cell model. The new parameter R modulates crossbridge formation as explained in the text. Cell model diagram modified from[1].

We chose the functions $f_1(R)$ and $f_2(R)$ to be of the form $f_i(R) = R^n$ based on the following considerations.

Considering the ordinary differential equation that describes the time evolution of weakly-bound crossbridges (W) reported above, in steady-state conditions, when $\frac{dW}{dt} = 0$, the relationship between the steady-state occupancies and transition rates will be

$$0 = k_{uw} * f_1(R) * U - k_{wu} * f_2(R) * W - k_{ws} * W$$

as no distortion ($\gamma_{wu}$) is present in steady-state, and therefore

$$\left(\frac{W}{U}\right)_{ss} = \frac{k_{uw} * f_1(R)}{k_{wu} * f_2(R) + k_{ws}}$$

The system will therefore reach a steady-state configuration determined by $f_1(R)$ and $f_2(R)$. In particular, depending on $R$, the ratio $\left(\frac{W}{U}\right)_{ss}$ will be shifted towards the U or the W state in line with the hypothesis that changing the availability of myosin heads that can interact with actin will affect how many crossbridges can enter in the pre-powerstroke state W. When $f_1(R) = f_2(R) = 1$, the original system is recovered (including the steady-state configuration). We therefore impose the constraint that $f_1(1) = f_2(1) = 1$, so that $R = 1$ represents the control condition where (DRX:SRX)=(DRX:SRX)$_{\text{control}}$. Note that $R \to 0$ as $\mathrm{DRX} \to 0$ (no interaction with actin), which implies $f_1(0) = 0$ (no crossbridges in pre-power state W). Moreover, $f_1(R)$ must be strictly monotonic increasing in $R$ to signify a gradual increased transition to the pre-powerstroke state W. Similarly, $R \to \infty$ ($R \gg 1$) as $\mathrm{SRX} \to 0$ (no crossbridges in super relaxed state, so all contribute to force production). Therefore, $f_2(R \gg 1) \to 0$, increasing the $\left(\frac{W}{U}\right)_{ss}$ ratio, where $f_2(R)$ must be strictly monotonic decreasing in $R$ to signify a gradual increased number of crossbridges remaining in the pre-power state W. Finally, both $f_1(R)$ and $f_2(R)$ must be strictly positive in the range of considered values of $R$ for compatibility conditions in the crossbridge cycling model (positive transition rates).

**Experimentally informed calibration of the human in-silico cellular model enables estimation of the relationship between myosin availability and Mavacamten concentration**

We calibrated the modified cellular model by determining the value of the $n$ parameter featured in $f_1(R)$ and $f_2(R)$ that achieved changes in contractile function by SRX:DRX modulation that are consistent with experimental data. Specifically, we considered the observed ~30% and ~70% decrease in maximal steady-state tension under 0.5μM and 1μM Mavacamten in human[9] and rat[10] myocardium, respectively, and the ~40% increase in contractility under an increase in myosin DRX of 17% due to the pathogenic HCM variant *MYH7*$^{\text{R403Q/+}}$, observed in human induced pluripotent stem cell-derived cardiomyocytes (hiPSC-CMs)[7], as calibration targets.

To achieve this, we conducted a sensitivity analysis in which we varied the parameter $n$ to generate a population of models. We considered the two scenarios of myosin inhibition (R < 1) and enhancement (R > 1) separately. We considered a positive $n$ for $f_1(R)$ and a negative $n$ for $f_2(R)$, since $f_1(R)$ and $f_2(R)$ must be strictly monotonic increasing and decreasing in $R$, respectively. We first determined the range of variation of $n$ by qualitatively evaluating the behaviours of $f_1(R)$ and $f_2(R)$ for selected $n$ values. Based on that, we decided to vary the $n$ parameter in the range [0.1, 0.5] with step size 0.1 for $f_1(R)$, and in [-3, -1] with step size 0.5 for $f_2(R)$ (Supplementary Fig. 2a). This is because beyond those ranges $f_1(R)$ and $f_2(R)$ would achieve extreme values that would lead to non-physiological model behaviours. We tested all 25 resulting combinations of $f_1(R)$ and $f_2(R)$ for R values in the physiological range [0.2, 6] with variable step size, and for each generated model we simulated the twitch and steady-state active tension (Supplementary Fig. 2b). We computed the twitch tension amplitude and maximal steady-state tension of each model, plotted them against corresponding R values (Supplementary Fig. 2c-e), and compared them to experimental evidence to determine the optimal parametrisation of the modified model.

From experimental evidence, a 40% increase in contractility was observed with a change in R value (R = 1.3), however there is no experimental evidence detailing the relationship between Mavacamten concentrations and R values. This necessitated that we determined this correspondence in silico, i.e. that we defined R values that, when used in our models, provide the same change in contractility in silico as it was observed with experimental data for 0.5 and 1μM Mavacamten concentrations.

As shown in Supplementary Fig. 2f, there exists a range of R values, one value for each model, for each of the two Mavacamten concentrations considered, that can satisfy that criterion, i.e. replicate the ~30% and ~70% decrease in maximal steady-state tension under 0.5 and 1μM Mavacamten observed experimentally. Therefore, we constructed an ensemble of dose-response curves where R values can be approximated from Mavacamten concentrations. We adopted the following modified Hill equation which resembles the dose-dependent effect of Mavacamten observed in experiments[10]:

$$R = \frac{a}{1 + \left(\frac{[Mava]}{IC50}\right)^n} + b.$$

We constructed a specific R-Mavacamten dose-response curve for each model in the population. To determine the parameters of each dose-response curve, we first considered the model's behaviour without Mavacamten exposure and at very high Mavacamten concentrations. Very high concentrations were here exemplified by 10μM, which is 4 to 8 times higher than the dose range of 1.3-2.6μM administered in clinical trials to achieve left ventricular outflow tract gradient below 30mmHg while preserving left ventricular ejection fraction in obstructive HCM patients[11,12]. Furthermore, experiments consistently showed that Mavacamten exerts the majority of its effects on healthy cells within the 1μM dose with an IC50 for cardiac myosin activity in the submicromolar range[10,13-15].

When there is no Mavacamten in the system, R should equal 1, whereas at 10μM Mavacamten, R should have approached the plateau of 0.2. This was considered here as the smallest possible value of R based on evidence suggesting that myosin DRX mostly varies between 20 and 60[7,8], as explained before. This leads to $b = 0.2$ and $a = 1 - b = 0.8$. This holds for all dose-response curves. Then, for each of the curves we estimated their specific IC50 and Hill coefficient $n$ using the MatLab fitting toolbox. Each fit was determined considering the following sets of points. 0μM Mavacamten should correspond to R = 1 and 10μM Mavacamten should correspond to R = 0.2. These two points are common to all dose-response curves, while 0.5 and 1μM Mavacamten have R values that are specific to each model. These are the estimated values and provide the expected tension under those two drug concentrations inferred from the experimental data. This procedure allowed us to parametrise each dose-response curve.

Finally, based on the above analyses, we were able to select one model in the population as our control baseline model. Specifically, we selected the model that (i) generated an R-

Mavacamten dose-response curve as close as possible to the average dose-response curve, and (ii) replicated the 40% increase in cardiac contractility expected for a myosin ratio of R = 1.3[7]. This was obtained for $f_1(R) = R^{0.3}$ and $f_2(R) = R^{-2}$.

This calibrated model provided a dose-response curve (Supplementary Fig. 2f, black solid line) that is close to the average dose-response curve computed considering all model parameterisations (Supplementary Fig. 2f, black dotted line). This is characterised by an IC50 of 0.7μM and a Hill coefficient of 2.3 (Supplementary Fig. 2f). This is in keeping with previous evidence showing a dose-dependent reduction of myosin activity by Mavacamten with a IC50 in the submicromolar range for cardiac myosin[10,15]. Consistently with its mathematical formulation, the chosen model also replicated baseline behaviour without Mavacamten exposure, and at high dose 10μM Mavacamten, which respectively can be simulated by R = 1 and R = 0.2 (Supplementary Fig. 2f). Mavacamten doses of 0.5μM and 1μM can be simulated by R = 0.7 and R = 0.4, respectively (Supplementary Fig. 2f). The resulting model captured a 40% contractility increase, quantified as simulated active tension amplitude, associated with R = 1.3 (Supplementary Fig. 2d, black line). This model parametrisation also led to the expected 30 and 70% reductions in maximal steady-state tension (Supplementary Fig. 2e, black line) for Mavacamten concentrations of 0.5μM and 1μM.

Simulation results from our sensitivity analysis showed that our parametrisation is robust and tolerates perturbations in the $n$ parameter, as the overall behaviour of all generated dose-response curves is consistent and does not significantly deviate from the chosen one (Supplementary Fig. 2f).

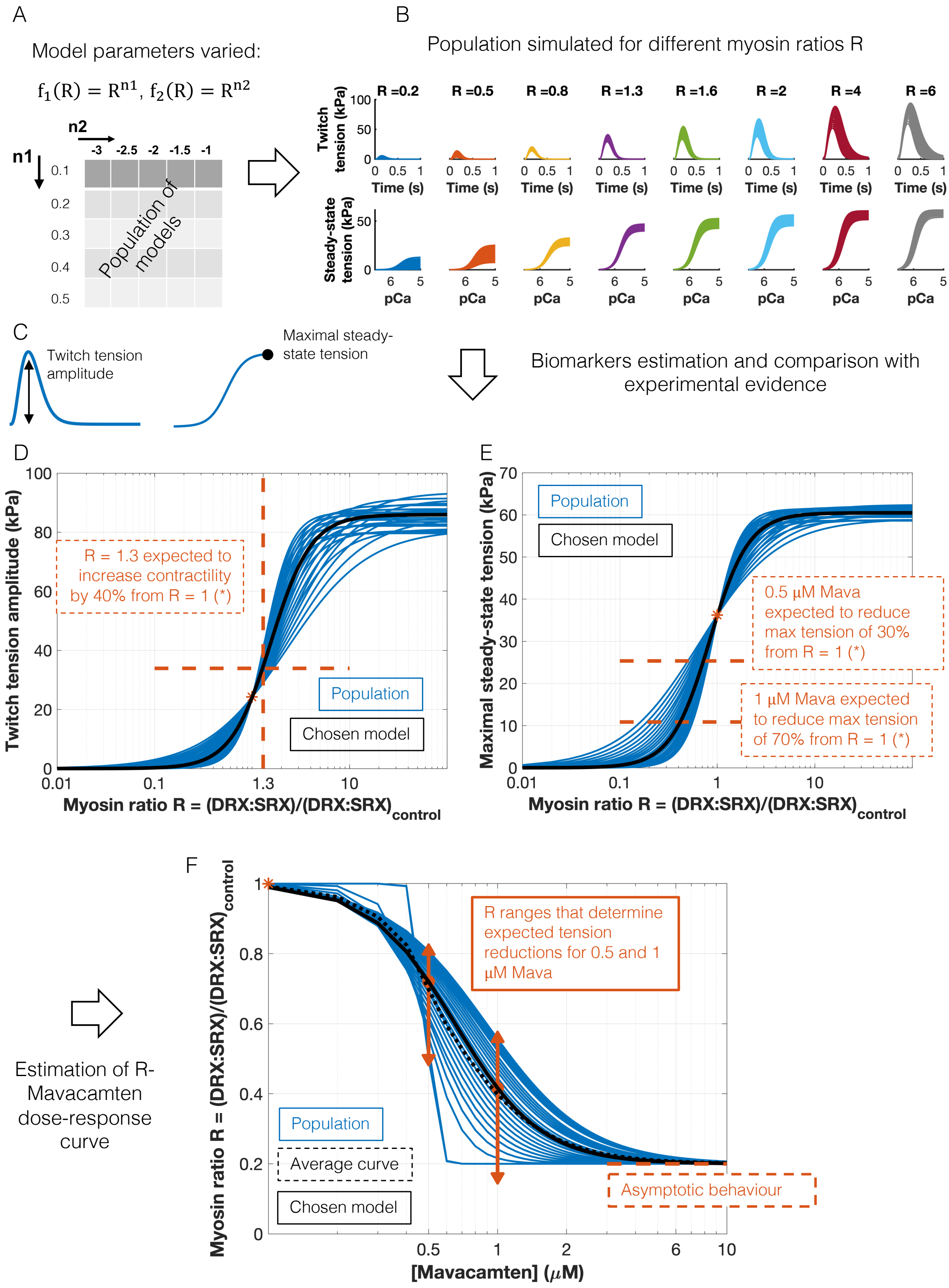


Supplementary Fig. 2. Calibration of the modified cellular electromechanical model and estimation of the R-Mavacamten dose-response curve. A: The $n$ parameter featured in $f_1(R)$

and $f_2(R)$ was varied as shown in the table, generating a population of models (see text for details). B: For each model in the population, we simulated the twitch and steady-state tension under different myosin ratios R. C-E: For each model we computed the twitch tension amplitude and maximal steady-state tension (C) and plotted them against the corresponding R values (D-E). The blue lines represent the fit through the computed biomarkers. The orange lines represent the constraints for model calibration derived from experimental evidence. Specifically, we considered the observed 40% increase in contractility under an increase in myosin DRX of 17% (R = 1.3) caused by a pathogenic HCM variant (D), and the observed 30 and 70% reduction in maximal steady-state tension under myosin modulation by 0.5$\mu$M and 1$\mu$M Mavacamten (E). The black solid line represents the calibrated model. F: Estimation of the R-Mavacamten dose-response curve for each model in the population (blue). The red symbols show the maximum, minimum, and median values of the myosin ratios R that determine, under 0.5$\mu$M and1 $\mu$M Mavacamten, the same amount of change in contractility in-silico with respect to experimental data. When there is no Mavacamten in the system, R is equal to 1 (red star), whereas for high Mavacamten concentrations, R reaches a plateau at 0.2 (asymptotic behaviour). The black dotted line represents the average dose-response curve whereas the solid black line represents the dose-response curve of the calibrated model.

**In-silico evaluation of preclinically relevant biomarkers of human cardiac contractility**

Two different protocols were simulated to obtain isometric twitch tension in human virtual intact cardiomyocytes and the steady-state tension-calcium relationship in human virtual skinned cardiomyocytes. Twitch tension is elicited in silico by the stimulus current that triggers an action potential through the excitation-contraction coupling process. The steady-state tension calcium curve is obtained by using fixed calcium values as input of the contractility model parametrised for skinned cardiomyocytes[5]. The tension produced by the cell quickly reaches a steady-state value that is saved and plotted with respect to the corresponding calcium value.

From these two protocols the relevant biomarkers were computed. The steady-state calcium-tension curve was fitted to a Hill equation. From this, maximal tension, calcium sensitivity of tension production, and Hill coefficient were estimated. From the twitch tension, time to 50% contraction, time to peak, time 50 and 90% decay, total duration, tension baseline and amplitude were computed. The model was paced at 1 Hz until the steady-state was reached, and then biomarkers were computed.

**In-silico modelling of myosin-based contribution to thin filament activation in human cardiomyocytes**

A crossbridge-dependent binding of calcium to cardiac troponin C has been observed experimentally[16,17] and also modelled computationally[18-21]. Building upon these findings, in this study we considered a direct modulation of thin filament calcium sensitivity based on the availability of myosin to form crossbridges and develop tension. Myosin availability is described in our model by the model parameter R which modulates the transition rates between the detached crossbridge state and the weakly bound one through the functions $\boldsymbol{f_1(R)}$ and $\boldsymbol{f_2(R)}$. To test our hypothesis of a myosin-based contribution to thin filament activation, we used the ratio $\frac{\boldsymbol{f_1(R)}}{\boldsymbol{f_2(R)}}$ to modulate thin filament activation, and specifically the Ca50 parameter. Of note, Ca50 represents the calcium concentration at half maximal thin filament activation and has $\boldsymbol{\mu}$M unit. Therefore, a larger Ca50 value means lower calcium sensitivity and vice versa. We constructed a Hill-type relationship between the $\frac{\boldsymbol{f_1(R)}}{\boldsymbol{f_2(R)}}$ ratio and the calcium sensitivity parameter Ca50 based on expected changes in tension relaxation due to myosin-based thin filament activation. Specifically, in the absence of an accurate experimental characterisation of the relationship between crossbridge cycling and thin filament activation, we considered data showing tension relaxation shortening under Mavacamten treatment[14], which reduces myosin availability, and data showing tension relaxation prolongation under myosin mutations that increase myosin availability[7], with the constraint that in control conditions, when $\frac{f_1(R)}{f_2(R)} = 1$, the Ca50 scaling should be 1. The resulting Hill curve, obtained with the Matlab Curve Fitting Toolbox, is illustrated in Supplementary Fig. 3 and the equation is reported below:

$$Ca50_{scaling} = \frac{0.3712}{1 + \left(\frac{f_1(R)/f_2(R)}{1.235}\right)^{4.21}} + 0.7368$$

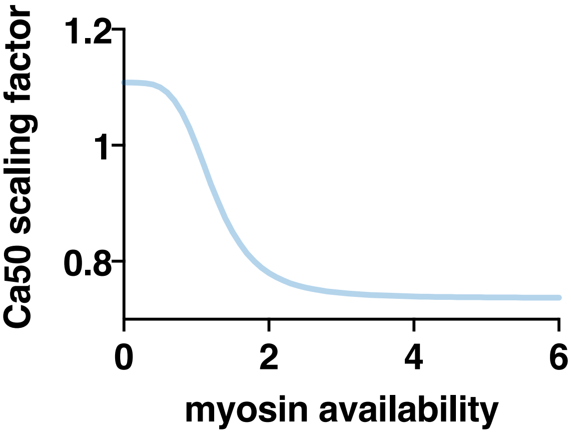


Supplementary Fig. 3. Modulation of calcium sensitivity of myofilaments by myosin availability implemented in the cellular model. The y axis reports the scaling factor of the parameter Ca50. A larger scaling factor determines an increase in Ca50 and therefore lower calcium sensitivity.

**Populations of in-silico human adult cardiomyocytes with thin filament troponin variants**

Different biophysical molecular defects have been reported as primary drivers of cellular dysfunction in HCM caused by thin filament mutations. The *TNNT2*$^{R92Q/+}$ mutation has been consistently reported to cause indirect calcium sensitisation of the myofilaments[22,23]. Recently it was also shown that a primary molecular insult for the observed hypercontractility and consequent disease pathogenesis in *TNNT2*$^{R92Q/+}$ is mutation-induced alterations in tropomyosin positioning[24]. We conducted a sensitivity analysis to determine the contribution of calcium sensitisation and abnormal tropomyosin positioning to altered cellular contractility. The reverse rate constant that defines $K_B$ (for tropomyosin positioning) in our baseline mechanical model and the parameter Ca50 that regulates calcium sensitivity of myofilaments were varied one at a time. $K_B$ represents the equilibrium constant between the blocked and closed tropomyosin states. Specifically, parameters were varied in the range $\pm$30% of their baseline value. We computed the absolute sensitivities of active tension and calcium transient biomarkers to changes in the model parameters as in[25]. From the sensitivity results on the relative contribution of tropomyosin positioning and calcium buffering in determining abnormal calcium and contractility under *TNNT2*$^{R92Q/+}$, an average mutant remodelling was selected (Ca50 x 0.7 and $K_B$ x 0.8, consistent with the reported increase in contractility under this mutation) and applied to the control population to generate a *TNNT2*$^{R92Q/+}$ population.

The *TNNI3*$^{R21C/+}$ mutation increases calcium binding of cardiac troponin and the affinity of cardiac troponin C for cardiac troponin I in solution, showing increases in the calcium sensitivity of myofibril tension development, and also prolonged early slow phase of relaxation[26]. Building upon these results we considered abnormal calcium-troponin binding as primary mutation-induced defect. We tested the contribution of changes in calcium sensitivity and dissociation rate from troponin to cellular contractility and calcium transient morphology by means of a sensitivity analysis. Specifically, the Ca50 parameter that describes the calcium sensitivity and the unbinding rate of calcium from troponin $k_{off}$ were considered. Ca50 was varied in the range $\pm$30% of its baseline value, and $k_{off}$ was varied in the range $\pm$50% of its baseline value. We computed the absolute sensitivities of active tension and calcium transient biomarkers to changes in the model parameters as in[25]. Similar to the troponin T variant, an average remodelling of calcium binding (Ca50 x 0.7) and dissociation ($k_{off}$ x 0.5) from troponin under *TNNI3*$^{R21C/+}$ was chosen so that it replicates the increase in contractility observed experimentally and was then applied to the control population to construct the *TNNI3*$^{R21C/+}$ population.

**In-silico evaluation of drug repurposing for the thin filament HCM phenotype correction**

In order to identify therapeutic targets for the phenotype resolution for the troponin variants, we also tested the effect of the following pharmacological strategies. We tested different levels of late sodium (Supplementary Fig. 4a,b) and L-type calcium (Supplementary Fig. 4c,d) currents block (20, 40, 60% block of current conductances) and different levels of SERCA upregulation (20, 50, 80% increase), in the presence and absence of 0.5µM Mavacamten (Supplementary Fig. 4e,f).

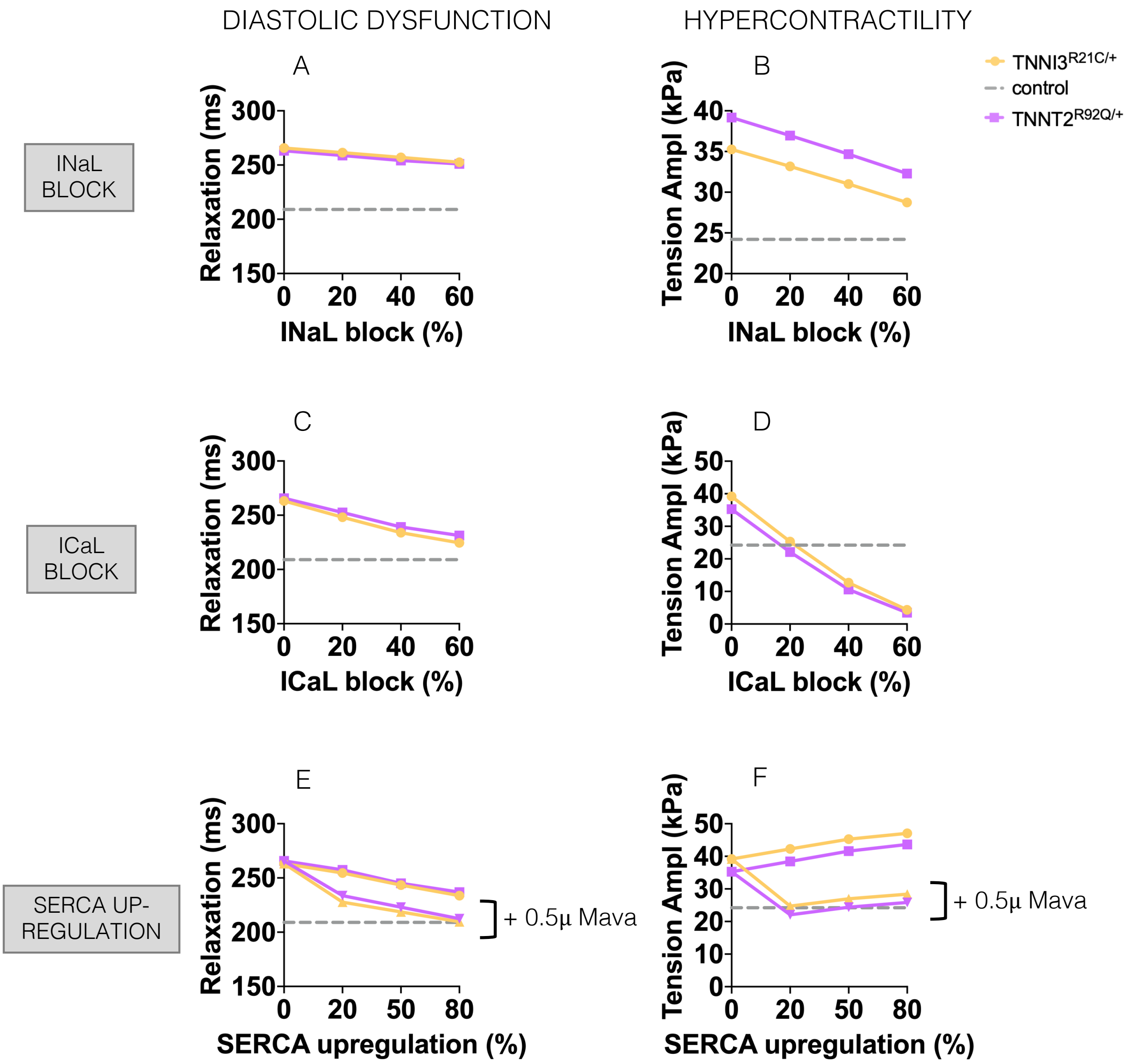


Supplementary Fig. 4. In-silico drug trials of different pharmacological strategies that can aid in the resolution of the thin filament HCM phenotype when Mavacamten action only is suboptimal. A&B: Effects of late sodium current block on tension relaxation (A) and amplitude (B). C&D: Effects of L-type calcium current block on tension relaxation (C) and amplitude (D).

E&F: Effects of SERCA upregulation, in the presence or absence of 0.5μM Mavacamtem (triangles), on tension relaxation (E) and amplitude (F).

**Calibration of Mavacamten cellular model with clinical data to enable simulation of clinically administered drug doses**

We conducted a recalibration of our cellular model of Mavacamten with the goal of, given a certain drug dose administered in clinical trials, determining what is the free plasma concentration to be used in the simulations to get appropriate left ventricular ejection fraction (LVEF) reductions. Available data from early phase clinical trials is reported in Supplementary Fig. 4a (data digitalised from[1]). Data was separated between healthy subjects and HCM patients. Only data from healthy subjects was used for calibration, as our whole-ventricular simulations only considered pre-symptomatic patients before development of obstructive HCM.

Before calibration, our in-silico trial results using preclinical dose concentrations reported a much stronger effect of Mavacamten on the LVEF than the clinical data (Supplementary Fig. 4b). To calibrate our cellular model of Mavacamten, we mapped simulated data onto clinical data to determine a correspondence between drug dose administered clinically ($DrugDose_{clinical}$) and simulated free plasma concentrations ($DrugDose_{simulated}$), as shown in Supplementary Fig. 4c. Clinical data can be written as

$$LVEF_{clinical} = a * DrugDose_{clinical} + b$$

where $LVEF_{clinical}$ represents the LVEF reduction and $DrugDose_{clinical}$ represents the corresponding plasma concentration measured from patients after drug administration. From fitting the clinical data of healthy volunteers reported in Supplementary Fig. 4a, $a$ and $b$ were estimated to be -0.0173 and 0.4732, respectively. Simulated data can be written as:

$$LVEF_{simulated} = \frac{c}{1 + \left(\frac{DrugDose_{simulated}}{d}\right)^{n}} + e$$

[1] https://www.streetinsider.com/SEC+Filings/Form+8-K+MyoKardia+Inc+For%3A+Sep+21/12057255.html

where $c, d, e, n$ were estimated to be 28.45, 117, -28.45, and 5.27, respectively (obtained from fitting the simulated data reported in Supplementary Fig. 4b). By writing $DrugDose_{simulated}$ as a function of $LVEF_{simulated}$ and considering $LVEF_{simulated} = LVEF_{clinical}$, we obtained $DrugDose_{mapped}$.

From $DrugDose_{mapped}$, calibrated myosin R values were then computed, to estimate the effect of the drug on myosin availability at the organ level, from the previously estimated equation

$$R_{clinical} = \frac{0.8}{1 + \left(\frac{DrugDose_{mapped}}{0.7}\right)^{2.3}} + 0.2$$

We computed the $R_{clinical}$ values corresponding to 500, 1000, and 1500ng/ml Mavacamten concentrations ($R_{clinical}$ = 0.86, 0.78, 0.64) and used them to conduct human virtual clinical trials.

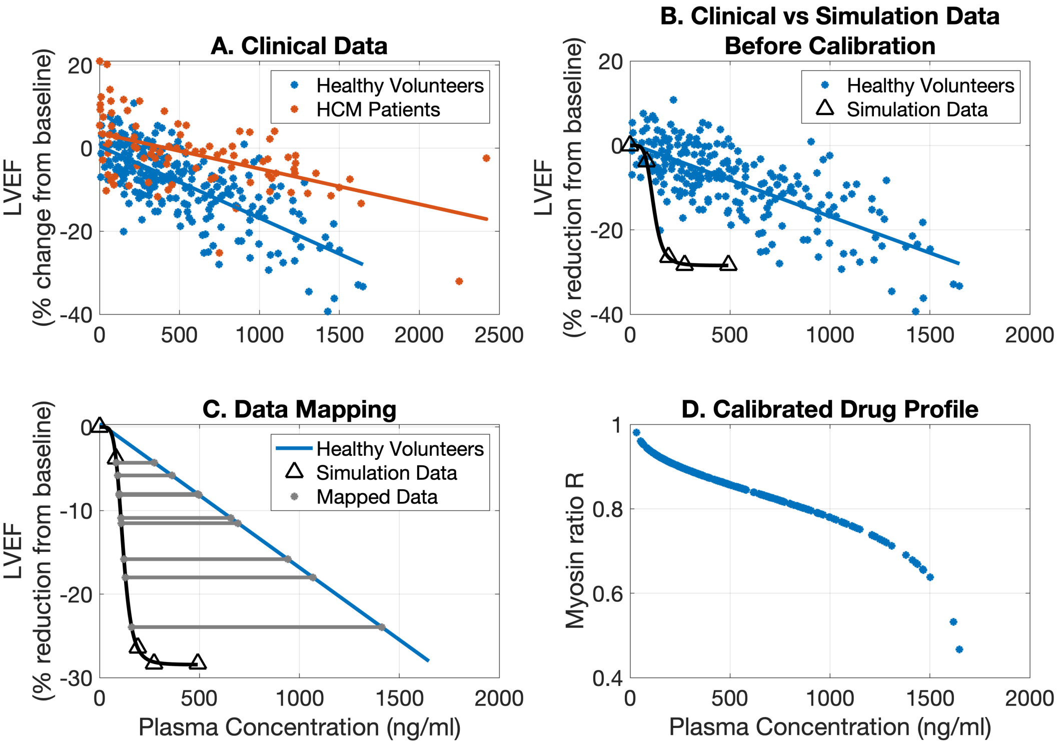


Supplementary Fig. 5. Calibration of our cellular model of Mavacamten to clinical data. A: The available clinical data is reported. B: Before calibration, our trials reported a much stronger effect on the LVEF than clinical data. C: Simulated data were mapped onto clinical data to determine a correspondence between drug dose administered clinically and simulated free plasma concentrations. D: The estimated free plasma concentrations were then used to recompute R values and determine the calibrated drug profile to be used in the human in-silico clinical trials.
